# Homologs of the Major Vault Protein defend bacteria against ribotoxins

**DOI:** 10.64898/2026.11.20.768214

**Authors:** Nathan P. Bullen, Lucia C. Holiday, Toni A. Nagy, Laurel K. Robbins, Aaron T. Whiteley

## Abstract

Bacteria use sophisticated mechanisms to defend themselves against antibiotics and bacteriophages, yet how they combat toxins deployed during interbacterial competition remains poorly understood. Bacteria use type VI secretion, bacteriocins, and contact dependent inhibition systems to deliver antibacterial toxins that inhibit growth by antagonizing essential processes. Here, we discover that *Pseudomonas aeruginosa* uses homologs of the eukaryotic major vault protein and a tRNA-like noncoding RNA to defend against RNA-damaging ribotoxins. Using an engineered pyocin delivery system, we showed that *P. aeruginosa* sensed ribotoxin damage and activated transcription of an RNA repair operon, which included *band7*, a homolog of the major vault protein. Genetic analyses demonstrated that *band7*-expressing bacteria were significantly more protected against growth inhibition by a ribotoxin that cleaved the anticodon loop of specific tRNA. Band7 was necessary but not sufficient, and defense also required a non-coding band7 RNA (bRNA). bRNA was cleaved by the ribotoxin, suggesting a mechanism for specificity and defense. Bioinformatic analyses revealed that *band7* and candidate *bRNA* are widely distributed within RNA repair systems across diverse bacteria. Our data suggest that bacteria use structures related to the eukaryotic vault complex to respond to RNA damage thereby providing a hypothesis for the function of the enigmatic human vault.

## Main text

Interbacterial antagonism is an inescapable aspect of bacterial life when bacteria compete for space and resources in complex environments ranging from the soil to sites of infection. A consequence of the relentless competition is the evolution of diverse antibacterial weapons that bacteria use to eliminate competitors. Among these are protein toxins that antagonize competing cells and kill by inhibiting essential processes, such as damaging cell envelope integrity, compromising the bacterial genome, or disrupting protein synthesis (*1*). Toxins are deployed through multiple mechanisms that each translocate protein cargo into a target cell. Type VI secretion systems (T6SSs) inject effectors directly into neighboring cells (*2*). Contact-dependent inhibition (CDI) systems deliver toxins following recognition of target-cell receptors (*3*). Protein bacteriocins are released into the extracellular environment and exploit cell-surface receptors and endogenous transport machinery to enter target cells (*4*). New toxins and delivery systems continue to be discovered, further expanding the known repertoire of weapons deployed during bacterial competition.

Survival during microbial competition is not just about deploying offensive strategies to kill competitors but also includes defensive strategies to survive threats. The threat of bacteriophages has driven the evolution of a staggering diversity of phage defense systems (*5*, *6*). Similarly, bacteria must also defend themselves against predatory bacteria (*7*, *8*). Perhaps the most famous defense strategy is antibiotic resistance, which evolved during intermicrobial competition long before clinical use of these compounds (*9*).

The diversity and abundance of antibacterial toxins deployed during microbial competition provides considerable selective pressure for bacteria to evolve mechanisms of defense against these threats. Toxin-producing bacteria neutralize their toxins using cognate immunity proteins that bind to the toxin with high affinity to block toxin activity and avoid self-intoxication (*10*). Closely related bacteria will also encode these immunity proteins to protect themselves, and one strategy of defense is to acquire immunity genes through horizontal gene transfer (*11*, *12*). However, less is known about how bacteria survive intoxication when they lack these cognate immunity proteins. Here, we used *Pseudomonas aeruginosa* as a model to investigate how bacteria survive the damage inflicted by antibacterial toxins.

### Engineering a toxin delivery platform

We searched the literature for antibacterial toxins produced by *P. aeruginosa* and identified 50 toxins with experimentally demonstrated antibacterial activity (Table S1). These toxins are deployed through three major delivery mechanisms: type VI secretion systems, contact-dependent inhibition, and S-type pyocins (soluble, pseudomonas-specific bacteriocins) (Fig. 1A) (*13*). Collectively, these toxins target essential cellular components that include the cell envelope (pore-forming toxins, phospholipases, and peptidoglycan hydrolases), the genome (DNases), coding and non-coding RNAs (RNases), and metabolites (NAD^+^ and ATP destroying-enzymes) (*1*). Toxins that cleave or modify tRNA, rRNA, and mRNA (ribotoxins) were the most prevalent in our survey (Fig. 1B). We hypothesized that *P. aeruginosa* must have evolved mechanisms to counter toxin-inflicted RNA damage and searched for mechanisms of ribotoxin defense.

**Figure 1.**
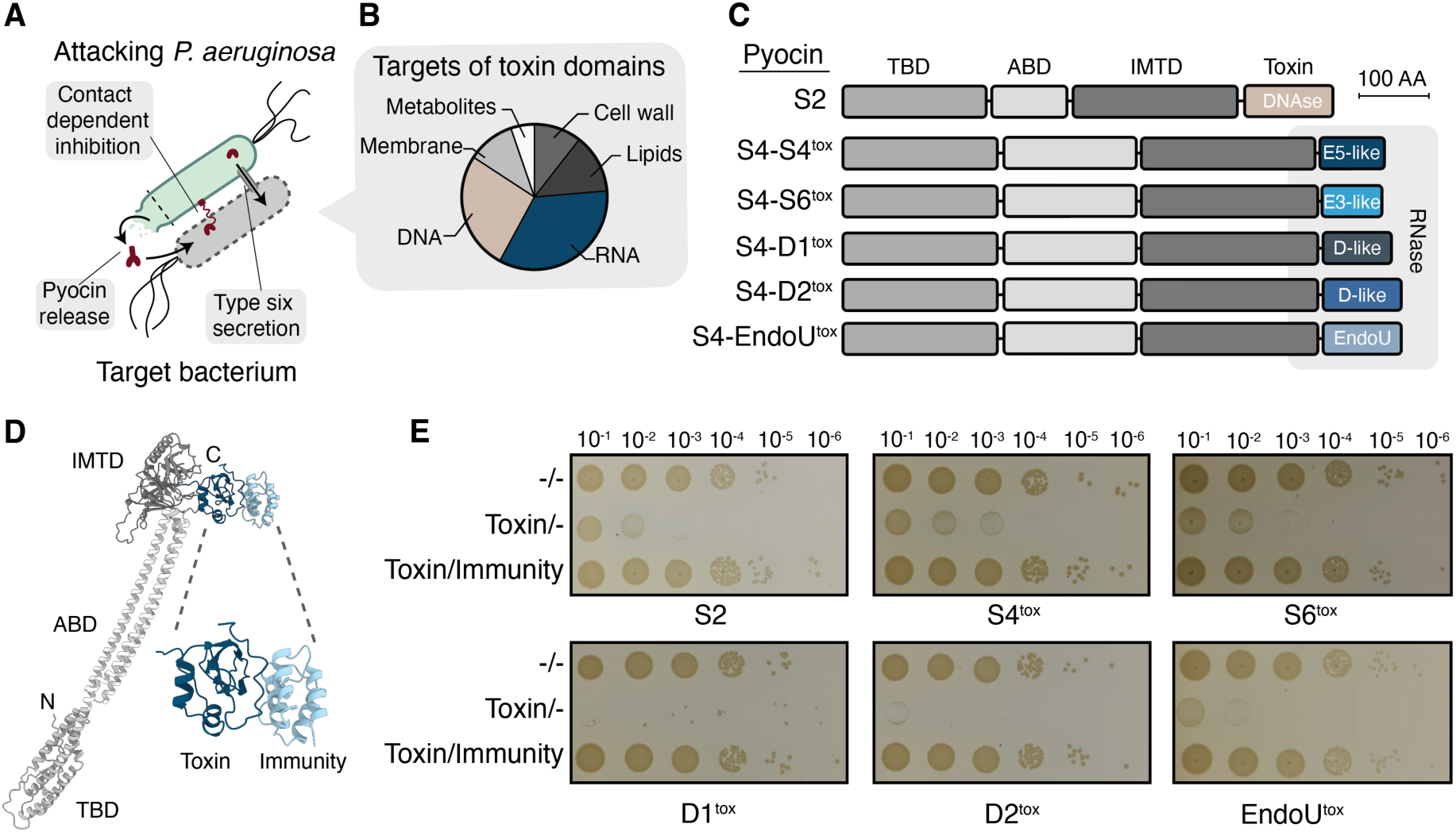
Pyocin S4 engineered into a RNAse delivery platform. **(A)** Schematic illustrating the major antibacterial toxin delivery mechanisms used by *P. aeruginosa*. Attacking *P. aeruginosa* cell is shown in green and the targeted bacterium in gray. **(B)** Distribution of cellular targets for experimentally characterized antibacterial toxin domains identified in our literature survey (Table S1). **(C)** Domain architectures of native pyocin S4 (S4-S4^tox^), engineered S4 ribotoxin chimeras, and pyocin S2 used in this study. Transporter-binding domain (TBD); Antigen-binding domain (ABD); Inner membrane translocation domain (IMTD) **(D)** AlphaFold-predicted structure of *P. aeruginosa* pyocin S4 in complex with its cognate immunity protein. The inset highlights the interaction between the toxin domain and the immunity protein. Domain colors and labels correspond to (C). **(E)** Toxicity assay showing tenfold serial dilutions of *P. aeruginosa* following exposure to the indicated toxins, spotted onto LB agar. For simplicity, the S4-prefix has been omitted and data are labelled according to toxin domains. -/-, no toxin exposure or immunity induction; Toxin/-, toxin exposure without immunity induction; Toxin/Immunity, toxin exposure with expression of the corresponding immunity protein induced with 250 µM IPTG.

We investigated diverse ribotoxins by engineering a system to deliver heterologous toxins into *P. aeruginosa* using an S-type pyocin as a delivery platform. Pyocin S4 uses an N-terminal receptor-binding and translocation domains to mediate uptake of a C-terminal toxin domain through the ferripyoverdine receptor FpvAI (*14*). The pyocin S4 toxin domain is related to colicin E5 and predicted to cleave tRNA (*15*, *16*). To repurpose the pyocin S4 delivery system, we substituted the native S4 C-terminal RNase domain with four alternative toxin domains representing distinct RNase families (Fig.1, C and D, Fig. S1A). These included the rRNase domain of pyocin S6 (S4-S6^tox^), which shares homology with colicin E3 and is predicted to cleave 16S rRNA; the tRNase domains of the contact-dependent inhibition toxins CdiA1 and CdiA2 from *P. aeruginosa* PA14 (S4-D1^tox^ and S4-D2^tox^), which shares homology with colicin D and is predicted to cleave tRNA anticodon loops; and the RNase domain of CdiA2 from PAO1 (S4-EndoU^tox^), which shares homology with the EndoU-family and whose cellular targets remain undefined (*16*, *17*). Together with native pyocin S4 (S4-S4^tox^), this yielded a panel of five purified ribotoxins with a range of targets (Fig. S1B). The DNA-targeting pyocin S2 was purified to use as a control for non-RNA damage (*18*).

We measured the impact of our chimeric pyocins on growth of *P. aeruginosa* by exposing bacteria to toxin and measuring colony formation. We selected strain PA14 for our analysis and, where necessary, deleted the endogenous copies of the toxin-antitoxin genes that would otherwise counter pyocin killing. Specifically, the toxin domains of S4-D1^tox^ and S4-D2^tox^ originate from PA14 and we deleted these toxins and cognate immunity proteins (Δ*cdiA-I*). A close homolog to pyocin S4 and its immunity protein were also found in PA14 and we deleted these genes (ΔPA14_13940-PA14-13950, Fig. S1A). Using the appropriate toxin-sensitive strains, we found all six pyocins potently inhibited bacterial growth (Fig. 1E). Next, we repeated the intoxication assay using strains heterologously expressing the immunity protein of each ribotoxin and found that each strain grew equally well to the no-toxin control (Fig. 1E). Immunity proteins inhibit their cognate toxins by specifically binding and occluding their active site, thereby providing an important control to demonstrate pyocin killing depended on the toxin domain. These data confirm that the observed growth inhibition for each chimeric pyocin was the result of the toxin domain’s activity.

### Ribotoxins activate an RNA damage response

We next investigated how *P. aeruginosa* responds to damage inflicted by ribotoxins by performing RNA sequencing of intoxicated cells. We focused our analysis on S4-EndoU^tox^ because examination of total cellular RNA from intoxicated cells revealed EndoU^tox^ induced the most grossly apparent RNA damage (Fig. 2A). After 5 minutes of intoxication with EndoU^tox^, the gene with the highest increase in abundance compared to non-intoxicated cells was PA14_60630 (86-fold), a gene of unknown function encoding a protein with a predicted Band7 domain, hereafter referred to as *band7* (Fig. 2B and S2A). Examination of the surrounding genomic region revealed that *band7* is encoded in an operon with three additional genes that were also strongly induced compared to cells not exposed to toxin: *rtcB*, encoding an RNA ligase (45-fold); *rlaP,* encoding a putative nucleotidyltransferase (12-fold); and *rtcA,* encoding an RNA 3′-phosphate cyclase (7.5-fold) (Fig. 2, B and C, Table S2). By 15 minutes of intoxication, *band7*, *rtcB*, *rlaP*, and *rtcA* had increased -590, 300-, 101-, and 56-fold, respectively (Fig. 2B, and Table S2).

**Figure 2.**
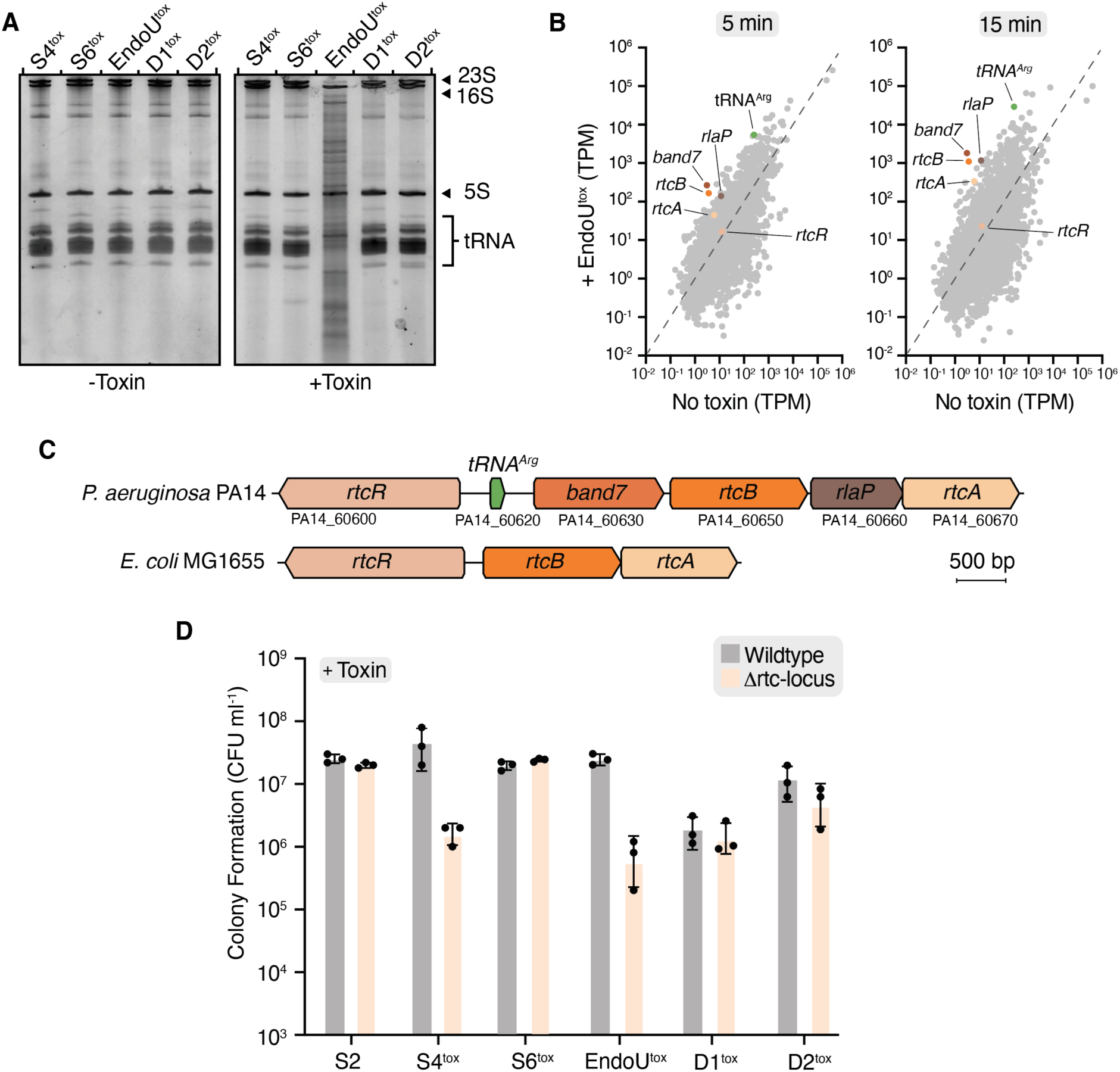
The *P. aeruginosa* PA14 Rtc operon protects against ribotoxins. **(A)** Urea-PAGE analysis of total cellular RNA extracted from *P. aeruginosa* pre-and post-15 min of exposure to the indicated ribotoxins. Major RNA species and cleavage products are indicated. **(B)** RNA-sequencing analysis of *P. aeruginosa* following 5 or 15 minutes of S4-EndoU^tox^ intoxication. Gene expression (transcripts per million, TPM) is plotted against untreated control cells. Genes within the *rtc* locus and the adjacent tRNA^Arg^ are highlighted. Dashed lines indicate equal expression between conditions. Colored data points correspond to (C). **(C)** Gene locus schematic of the *P. aeruginosa* and *E.coli rtc* operons. Locus tags for the *P. aeruginosa* operon are indicated under each gene. **(D)** Quantification of colony-forming units (CFU ml^-1^) of wildtype and Δrtc-locus *P. aeruginosa* PA14 following exposure to the indicated toxins. Data are the mean ± standard error of the mean (SEM) of three biological replicates (n = 3), with individual replicates shown as points.

Together, *band7*, *rtcB*, *rlaP*, and *rtcA* constitute the Rtc operon of *P. aeruginosa* PA14, which is regulated by the divergently encoded σ^54^-dependent transcriptional activator RtcR (Fig. 2C). RNA repair systems (Rtc operons and RtcR) are widespread among bacteria and their physiological roles remain poorly understood (*19*, *20*). Although these systems are named for RNA repair, their functions can also include RNA damage responses that mitigate, rather than heal, damage (*21*, *22*). In a comprehensive analysis of bacterial RNA repair systems, Burroughs and Aravind found that bacteria encode a broad-range of diverse genes within their Rtc operons and neighboring genomic regions (*23*). The canonical Rtc operon in *E. coli* encodes only the minimal *rtcB* and *rtcA* genes divergently transcribed from *rtcR* (Fig. 2C). However, many bacteria encode additional components with known or predicted RNA repair functions. *P. aeruginosa* has broadened its Rtc operon to include *band7* and *rlaP*. Rtc-related RNA repair loci show patterns of gene shuffling and horizontal transfer that are consistent with the rapidly evolving systems involved in microbial conflict (*23*). Given the strong induction of the PA14 Rtc operon following ribotoxin treatment, we hypothesized these genes function as a defense pathway to counter antibacterial ribotoxins.

We deleted the PA14 Rtc locus (Δrtc-locus), challenged this mutant with our toxin panel, and measured colony formation. We recovered over 50-fold fewer colony forming units (CFU) of the Δrtc-locus mutant compared to wild-type when treated with either S4-EndoU^tox^ or S4-S4^tox^, but saw no change in sensitivity of Δrtc-locus to the other four pyocins (Fig. 2D). Importantly, the Δrtc-locus mutant had no growth defect in the absence of toxin (Fig. S2B).

We next challenged the Δrtc-locus strain with diverse biotic and abiotic stressors, including UV irradiation, temperature stress, redox stress, bacteriophage infection, and varied antibiotics, to further examine the specificity of this defense response (see Methods). For each stressor, the Δrtc-locus mutant was as equivalently sensitive as wild-type (Fig. S2, C, D, E and F). Together, our findings suggest that the Rtc operon functions as a specialized defense pathway against specific ribotoxin-mediated RNA damage.

### Rtc operon genes provide ribotoxin defense

We next identified the genes within the Rtc operon that were responsible for defense against pyocin S4-S4^tox^ and S4-EndoU^tox^ by generating in-frame deletions of each gene. *band7* was required for defense against S4-EndoU^tox^, *rtcB* was required for defense against S4-S4^tox^*, rlaP* was dispensable for both toxins, and *rtcA* was required for protection against both toxins (Fig. 3, A and B). The degree of ribotoxin defense was not equivalent for each individual rtc gene. Specifically, the Δ*band7* and Δ*rtcB* mutants were similarly sensitive to intoxication as the Δrtc-locus strain for S4-EndoU^tox^ and S4-S4^tox^, respectively. In contrast, deletion of *rtcA* resulted in an intermediate sensitivity for both toxins.

**Figure 3.**
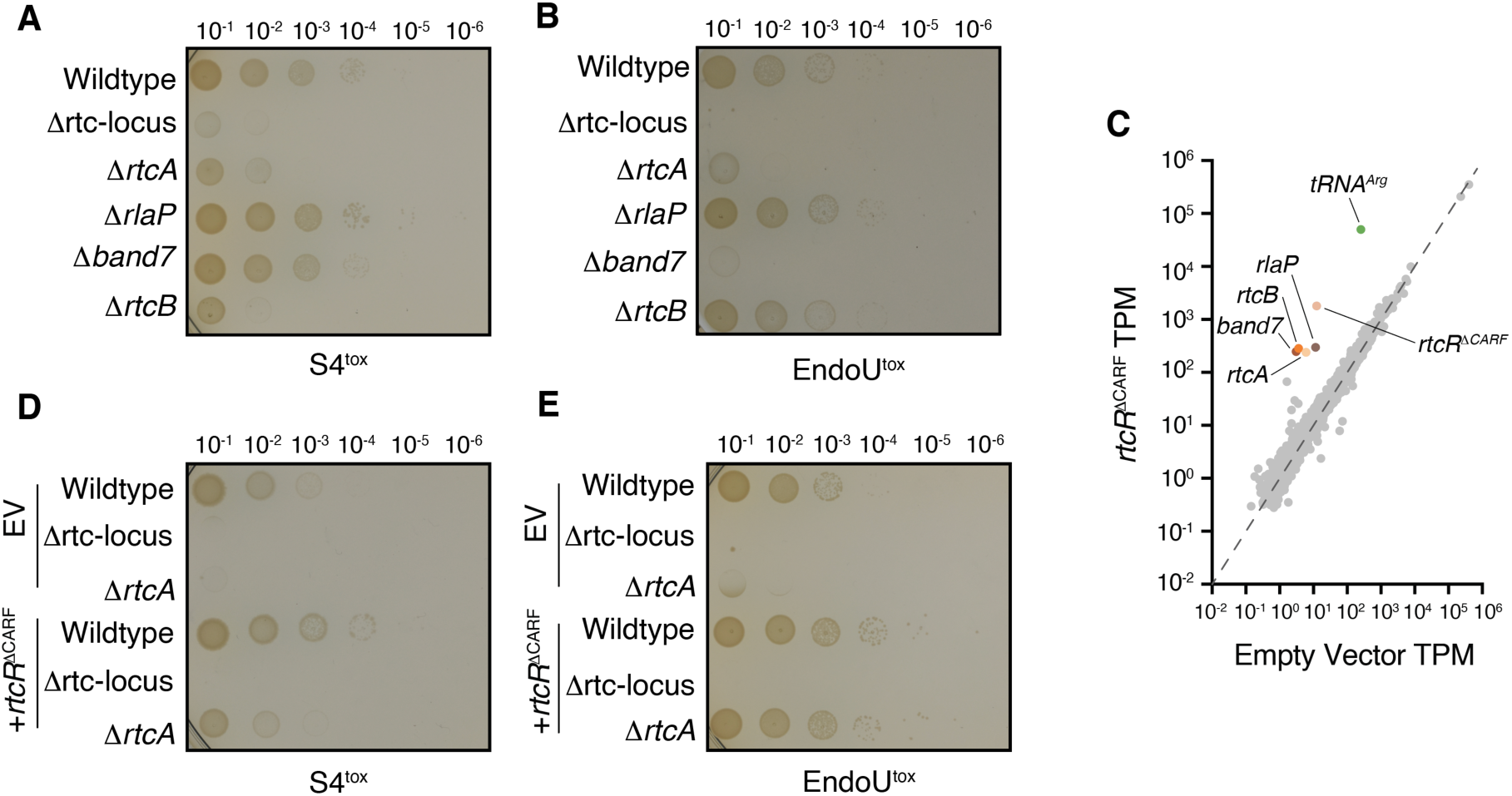
Rtc locus components required for distinct ribotoxins. **(A and B)** Toxicity assays showing tenfold serial dilutions of the indicated *P. aeruginosa* mutants following exposure to S4-S4^tox^ (A) or S4-EndoU^tox^ (B), spotted onto LB agar. **(C)** RNA-sequencing analysis of *P. aeruginosa* 20 minutes after induction of *rtcR*^ΔCARF^ with 250 µM IPTG. Gene expression (transcripts per million, TPM) is plotted against an empty vector containing control strain. Genes within the Rtc locus and the adjacent tRNA^Arg^ are highlighted. Dashed lines indicate equal expression between conditions. **(D and E)** Toxicity assays showing tenfold serial dilutions of the indicated *P. aeruginosa* strains expressing *rtcR*^ΔCARF^ or carrying an empty vector control (EV), following exposure to S4-S4^tox^ (D) or S4-EndoU^tox^ (E). Cells were spotted onto LB agar supplemented with 250 μM IPTG.

RtcR activates transcription of the divergently transcribed Rtc operon upon binding of damaged RNA fragments (*24*, *25*). The RtcR N-terminal CARF domain recognizes RNA damage by binding 2′,3′-cyclic phosphate termini, which frees the AAA+ ATPase domain to interact with σ^54^ and leads to increased target gene expression (*19*, *26*). One of the sources of cyclic 2′,3′-cyclic phosphate termini is RtcA, which converts RNA 3′-phosphate ends to 2′,3′-cyclic phosphates (*20*). Our transcriptomics suggested RtcR was activated following ribotoxin damage and we hypothesized that RtcA might help to potentiate this response by increasing the abundance of 2′,3′-cyclic phosphate termini, thereby explaining the Δ*rtcA* intermediate phenotype for S4-EndoU^tox^ and S4-S4^tox^. To test this hypothesis, we measured ribotoxin sensitivity during heterologous expression of a constitutively active RtcR variant deleted for its N-terminal CARF domain (*rtcR^ΔCARF^*). RNA-sequencing analysis confirmed that *rtcR^ΔCARF^* induced expression of the Rtc operon in the absence of intoxication (Fig. 3C). Further, expression of constitutively active *rtcR^ΔCARF^* rescued the sensitivity of Δ*rtcA* strains (Fig. 3, D and E). These data suggest that the principal function of RtcA in these conditions is amplifying expression of the Rtc operon through production of RtcR-activating ligands.

Unlike in the Δ*rtcA* mutant, expression of constitutively active *rtcR^ΔCARF^* failed to rescue the sensitivity of the Δ*rtcB* strain to S4-S4^tox^, indicating that RtcB is directly responsible for defense against this ribotoxin (Fig. S3A). RtcB is an RNA ligase that joins cleaved RNA ends bearing 3′-phosphate or 2′,3′-cyclic phosphate with 5′-hydroxyl termini (*27*, *28*). Colicin E5-family ribotoxins, including pyocin S4^tox^, cleave tRNAs within their anticodon loops, preventing their use in translation (*29*). Previous studies have shown that *E. coli* RtcB can directly repair tRNAs cleaved within their anticodon loops *in vitro* (*27*). Our data, suggest that RtcB protects against S4-S4^tox^ through ligation to directly repair pyocin-cleaved tRNAs.

Heterologous expression of constitutively active *rtcR^ΔCARF^* also did not affect the sensitivity of the Δ*band7* mutant to S4-EndoU^tox^, implying that Band7 directly mediates defense against this toxin (Fig S3B). Band7 domain-containing proteins have not been previously shown to contribute to RNA damage defense, prompting us to further investigate their molecular function.

### A homolog of major vault protein and a non-coding RNA are required for defense

Band7 proteins belong to the widespread stomatin, prohibitin, flotillin, HflK/C (SPFH) superfamily, whose members are often associated with membrane organization and the formation of large oligomeric assemblies (*30*, *31*). A previous bioinformatic analysis of bacterial RNA repair operons noted that the subset of bacterial Band7 proteins in these loci share distant homology with the eukaryotic major vault protein (MVP), and raised the possibility that these proteins form vault-like assemblies (*23*). Vaults are massive, barrel-shaped ribonucleoprotein complexes composed primarily of MVP, and are of largely unknown function (*32*, *33*). We examined the predicted structure of PA14 Band7 and found striking resemblance to MVP, including a similar arrangement of N-terminal repeat domains, a shared Band7/SPFH shoulder domain, and a C-terminal capping helix (Fig. 4A). Modeling the oligomeric complex of 12 Band7 monomers revealed a vault like complex comparable in shape to the human vault (Fig. 4B).

**Figure 4.**
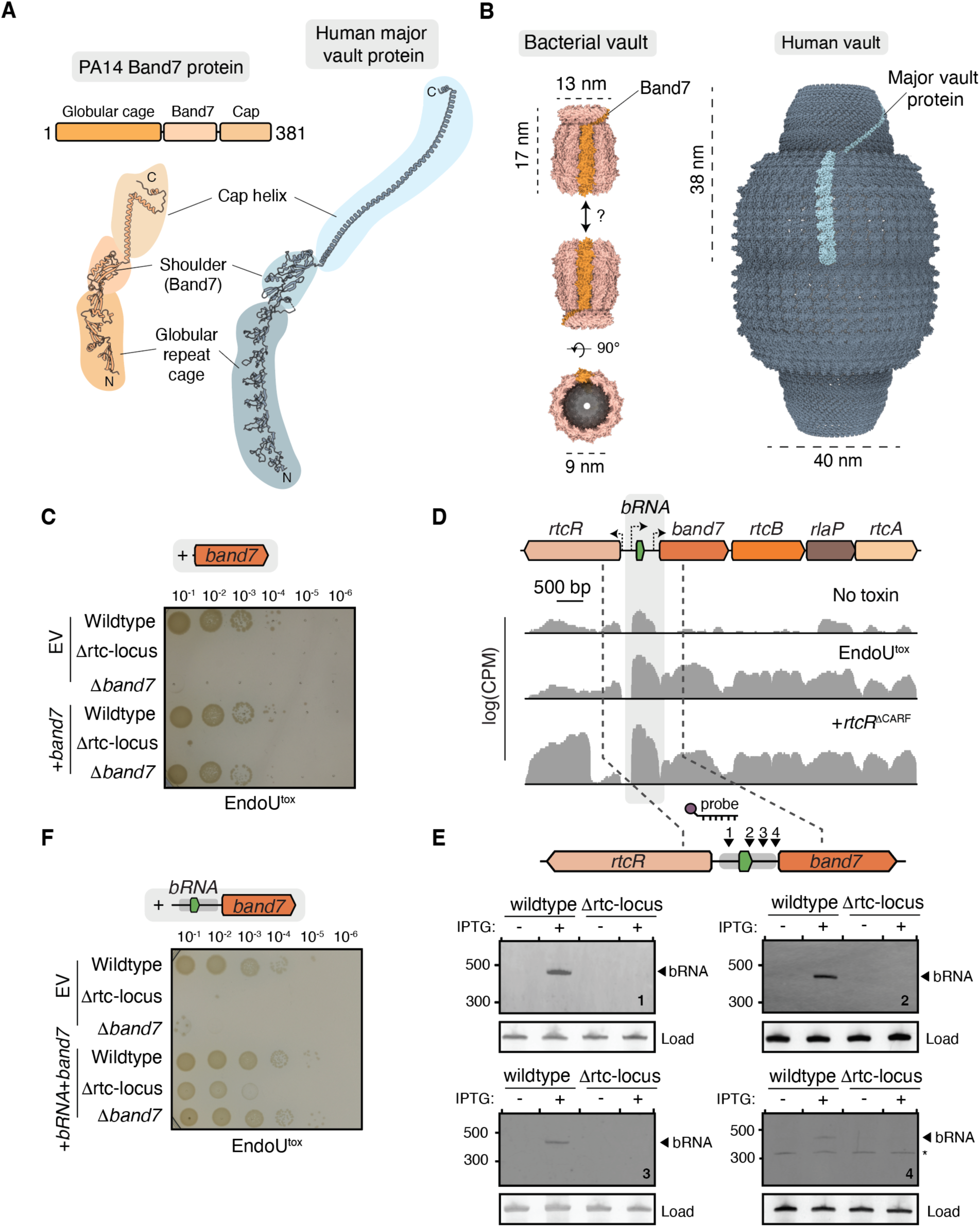
Band7 and a tRNA-like noncoding RNA confer protection against ribotoxin-mediated damage. **(A)** Domain architecture and AlphaFold-predicted structural model of *P. aeruginosa* Band7 protein compared to the structure of the human major vault protein (PDB 9BW5). Corresponding structural features are indicated. **(B)** Predicted oligomeric assembly of the *P. aeruginosa* Band7 modelled as a 12-mer compared with the human vault complex (PDB 9BW5). A protomer in each structure is highlighted. **(C)** Tenfold serial dilutions of wildtype *P. aeruginosa*, Δrtc-locus, and Δ*band7* strains carrying either an empty vector (EV) or a plasmid expressing *band7*, following intoxication with S4-EndoU^tox^. Cells were spotted onto LB agar supplemented with 250 μM IPTG. **(D)** RNA-sequencing coverage across the *P. aeruginosa* Rtc operon in untreated cells, cells intoxicated with S4-EndoU^tox^, and cells expressing RtcR lacking its N-terminal CARF domain (*rtcR*^ΔCARF^). Coverage is shown as log-transformed counts per million (CPM). The annotated tRNA-like region upstream of *band7*, subsequently identified as part of bRNA, is highlighted. **(E)** Northern blot analysis of bRNA expression using four probes targeting the indicated regions of the transcript. RNA was isolated from wildtype *P. aeruginosa* and the Δrtc-locus mutant grown in the presence or absence of IPTG. Arrowheads indicate the approximately 450-nucleotide bRNA transcript. A background band for probe 4 is indicated by *.The 5S rRNA imaged prior to electrophoretic transfer of RNA to the membrane is used as loading control. **(F)** Tenfold serial dilutions of wildtype PA14, Δrtc-locus, and Δ*band7* strains carrying either an empty vector (EV) or a plasmid expressing *bRNA* and *band7*, following intoxication with S4-EndoU^tox^. Cells were spotted onto LB agar supplemented with 250 μM IPTG.

The *band7* gene was required for defense against S4-EndoU^tox^ and we next attempted to show that *band7* expression was sufficient to restore the Δrtc-locus mutant sensitivity to toxin. Unexpectedly, expression of *band7* did not restore defense in the Δrtc-locus strain. This was unlikely to be technical failing because complementation of the Δ*band7* mutant with the same *band7* expression vector restored protection against S4-EndoU^tox^ (Fig. 4C). These findings suggested that an additional component within the rtc-locus was required for defense, but our other data showed this component was not *rtcB, rlaP,* or *rtcA*. To discover the unknown component, we returned to our *rtcR^ΔCARF^*RNA-seq results and closely examined transcription across the Rtc locus. Transcription appeared to originate from three locations: from the promoter of *rtcR*, from the promoter of the Rtc operon, and in the intergenic region between rtcR and the Rtc operon (Fig. 4D). Notably, the intergenic region transcript included an annotated arginyl-tRNA gene.

Northern blot analysis using probes targeting this region in cells expressing *rtcR^ΔCARF^*revealed a distinct RNA species of approximately 450 nucleotides, substantially longer than the 75-90 nucleotides expected for a mature arginyl-tRNA (Fig. 4E). Based on RNA-sequencing coverage and the estimated transcript size, this RNA’s transcription start site appeared to be approximately 140 nucleotides upstream of the annotated tRNA, with transcription terminating roughly 50 nucleotides upstream of the *band7* coding sequence. We renamed this non-coding RNA *<u>b</u>and7*-associated <u>RNA</u> (bRNA). To determine whether bRNA contributes to defense, we heterologously expressed both *bRNA* and *band7* in Δ*band7* and Δrtc-locus mutants and challenged these strains with S4-EndoU^tox^. This construct restored protection to Δ*band7* and Δrtc-operon, unlike expression of *band7* alone (Fig. 4F). Introduction of a premature stop codon into the *band7* coding sequence abolished protection, demonstrating that both the Band7 protein and upstream noncoding RNA are required for defense (Fig. S4). Together, these findings reveal that crucial elements of the Rtc operon ribotoxin defense system include a vault-like Band7 protein and an associated noncoding bRNA.

### S4-EndoUtox cleaves tRNAs near their anticodon loops

Antibacterial ribotoxins kill cells by cleaving essential RNAs, particularly tRNAs and rRNAs (*1*). Unlike the other ribotoxins in our panel, sequence and structural predictions provided little insight into the potential RNA targets of S4-EndoU^tox^. We investigated how this toxin inhibits bacterial growth by performing northern blot analysis of RNA isolated from intoxicated cells using probes targeting six distinct tRNA species, 16S rRNA, and 23S rRNA. Probes targeting the variable loop regions of tRNAs Arg^ACG^, Arg^CCU^, Arg^CCG^, and Glu^UUC^ revealed distinct cleavage products of approximately 35-40 nucleotides (Fig. 5A, B and S5A). The size of these 3′ fragments was consistent with cleavage of each tRNA within its anticodon loop. To further investigate the position of cleavage, we repeated the analysis of Arg^ACG^ cleavage using a probe targeting the region 5′ of the anticodon loop (Fig 5B and C). This revealed a similarly sized cleavage product, consistent with the accumulation of the 5′ half of the cleaved tRNA. In contrast, we did not detect cleavage products for Ser^GCU^ or Val^UAC^, suggesting that EndoU^tox^ targets a subset of tRNA species rather than cleaving tRNAs indiscriminately (Fig. S5B).

**Figure 5.**
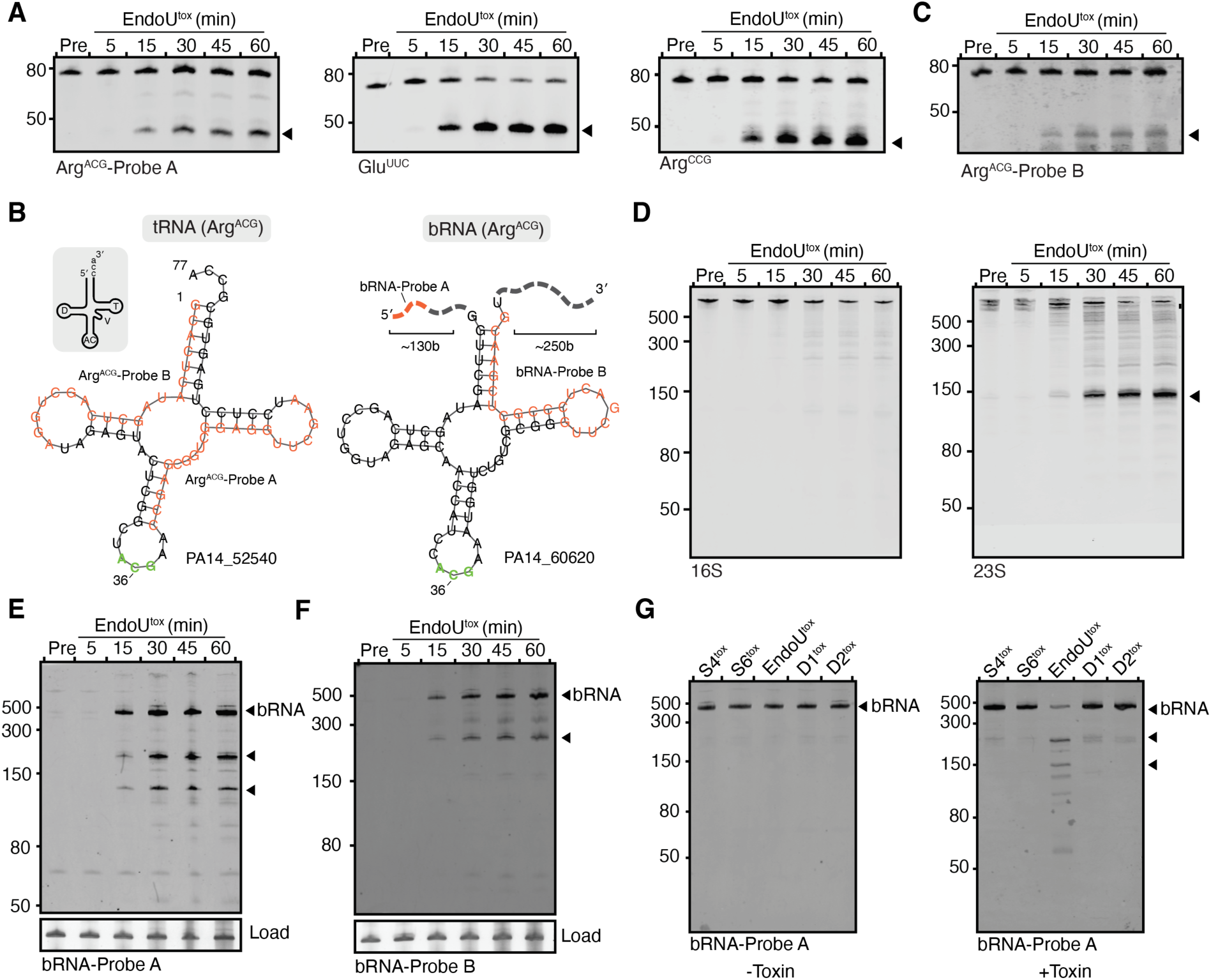
EndoU^tox^ selectively cleaves tRNAs, 23S rRNA, and the Band7-associated bRNA. **(A)** Northern blot analysis of tRNA cleavage following intoxication with S4-EndoUtox. RNA was isolated from PA14 cells before intoxication (Pre) or after 5, 15, 30, 45, and 60 min of intoxication. Probes targeting regions 3′ (Probe A) or 5′ (Probe B) of the anticodon loop of tRNA Arg^ACG^ are indicated and correspond to labels in (C). Arrowheads indicate major cleavage products. **(B)** Northern blot analysis of 16S and 23S rRNA following S4-EndoUtox intoxication. RNA was isolated from *P. aeruginosa* before intoxication (Pre) or at the indicated times post-intoxication. An approximately 150-nucleotide cleavage product of 23S rRNA is indicated by the arrowhead. **(C)** Secondary structure prediction depicting cloverleaf structure of tRNA Arg^ACG^ and the tRNA-like region of bRNA. Nucleotides are numbered according to their positions within respective tRNA-like sequences. The positions of probes used to target 5′ (Probe A) and 3′ (Probe B) of the anticodon region for each molecule are indicated in orange. The schematic in the top left depicts conserved features of the tRNA cloverleaf, including the D-loop (D), the anticodon loop (AC), the T-loop (T), and the CCA acceptor stem. **(D)** Northern blot analysis of bRNA cleavage following intoxication with S4-EndoUtox. RNA was isolated from *P. aeruginosa* before intoxication (Pre) or at the indicated times post-intoxication. Probes targeting regions 3′ (Probe A) or 5′ (Probe B) of the anticodon loop of bRNA Arg^ACG^ are indicated and correspond to labels in (C). Arrowheads indicate major cleavage products. **(E)** Northern blot analysis of bRNA cleavage following intoxication with the indicated ribotoxins. RNA was isolated from *P. aeruginosa* before intoxication (Pre) or at the indicated times post-intoxication. Probes used are as in (D). Arrowheads indicate major cleavage products.

Northern blot analysis of 23S rRNA also revealed a prominent cleavage product of approximately 150 nucleotides (Fig. 5D). This observation was consistent with analysis of total cellular RNA, which showed a near complete loss of full-length 23S rRNA following intoxication with S4-EndoU^tox^ (Fig. 2A). In contrast, we detected no comparable cleavage of 16S rRNA (Fig. 5D). Together, these findings suggest that S4-EndoU^tox^ inhibits bacterial growth by damaging multiple components of the translation machinery, including cleavage of a subset of tRNAs near their anticodon loops and degradation of 23S rRNA.

### bRNA is cleaved during S4-EndoUtox intoxication

bRNA encompasses an internal 74-nucleotide sequence annotated in the PA14 genome as an arginyl-tRNA with an ACG anticodon. Secondary structure prediction confirmed this region adopts a canonical tRNA cloverleaf structure (Fig. 5B). Sequence alignment with the five other arginine tRNA genes annotated in the PA14 genome revealed that this region retains conserved sequence motifs characteristic of the tRNA (i.e. D- and TΨC-loops) but lacks a genetically encoded CCA at the 3′-terminus, which is found on the other five arginyl-tRNA (Fig. S5C). RNA sequencing failed to detect post-transcriptional addition of a CCA 3′-end and our northern blot analysis did not identify bRNA at the expected size of mature tRNA. This suggests that bRNA is not a mature tRNA, unlikely to be amino acylated, and probably not participating in translation.

Given the similarity of bRNA to tRNA, we investigated whether bRNA is cleaved during S4-EndoU^tox^ intoxication. Northern blot analysis using a probe targeting the 5′ region upstream of the tRNA-like features revealed two prominent cleavage products following treatment with S4-EndoU^tox^. The first migrated between the 150- and 300-nucleotide markers and the second just below the 150-nucleotide marker (Fig. 5E). A second probe targeting the region 3′ of the tRNA-like features revealed a prominent cleavage product migrating below the 300-nucleotide marker (Fig. 5F). These sizes are consistent with cleavage near the predicted anticodon-loop like feature of bRNA, similarly to the observed cleavage of tRNA Arg^ACG^.

We next determined whether bRNA cleavage was associated with coordinating defense against S4-EndoU^tox^ intoxication. If bRNA cleavage is linked to this protective response, we reasoned cleavage should only occur specifically during intoxication with S4-EndoU^tox^ and not with ribotoxins from our panel for which Band7-bRNA did not provide protection. Northern blot analysis following intoxication with each of the five ribotoxins showed that only S4-EndoU^tox^ produced detectable bRNA cleavage (Fig. 5G). These findings establish that bRNA undergoes selective cleavage during S4-EndoU^tox^ intoxication, suggesting the tRNA-like features of bRNA may be important for sensing and responding to specific tRNA-targeting toxins.

### Bacterial vault-like proteins are widespread and associate with RNA repair systems

We next sought to determine how widely vault-like Band7 proteins are distributed among bacteria and whether their association with RNA repair systems is conserved. Initial sequence similarity searches using PA14 Band7 as a query returned many non-vault-like members of the SPFH superfamily, such as HflK/C homologs with N-terminal transmembrane helices rather than the globular N-terminal repeats characteristic of the major vault protein. To overcome this limitation, we used PSI-BLAST to curate a list of 19 Band7 proteins from distinct bacterial species that were confirmed to adopt N-terminal globular repeat domains and resemble PA14 Band7 and MVP. We then used these 19 sequences to construct a profile Hidden Markov model (HMM) designed to capture sequence features conserved among vault-like Band7 proteins while distinguishing them from other SPFH family members. Searching 88.8 million proteins from bacterial and archaeal RefSeq genomes assemblies with this model identified 2,026 high-confidence homologs.

Phylogenetic analysis of the Band7 homologs identified in our search revealed that vault-like Band7 proteins are distributed across most major bacterial phyla, including Pseudomonadota, Actinomycetota, Bacillota, and Bacteroidota (Fig. 6A). Structural modeling of representative proteins revealed conservation of the overall vault architecture, with predicted complexes varying in size and shape according to the number of N-terminal repeat domains and the length of the cap helix (Fig. 6B and C). Examination of the genomic neighborhoods surrounding these genes revealed that many are associated with Rtc RNA repair systems: adjacent to transcriptional regulators and enzymes with predicted RNA repair functions, including homologs of RtcB, RtcA, and RlaP (Fig. 6B and C). Intriguingly, many *band7* genes were also encoded immediately downstream of annotated tRNA-like sequences, resembling the genomic arrangement of *bRNA* and *band7* observed in PA14. These sequences were identified across distantly related bacterial lineages and corresponded to different predicted tRNA identities, suggesting that the association between vault-like Band7 proteins and tRNA-like bRNAs is a conserved feature of these systems. Together, these analyses reveal that vault-like Band7 proteins are phylogenetically widespread and retain a conserved association with RNA repair machinery, suggesting an ancient and conserved role in bacterial ribotoxin defense.

**Figure 6.**
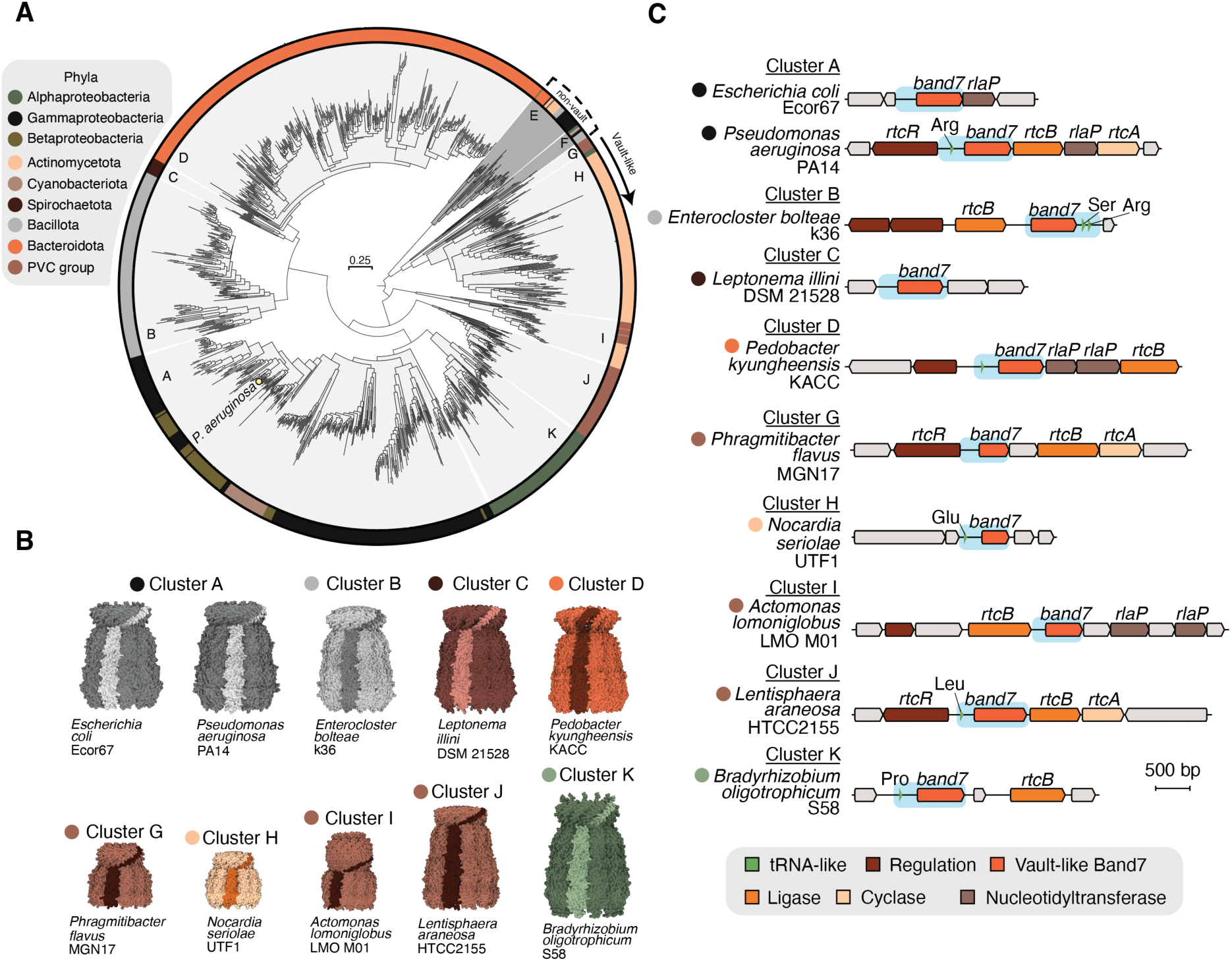
Vault-like Band7 proteins are widespread and frequently associate with RNA repair systems. **(A)** Maximum-likelihood phylogeny of 1,313 vault-like Band7 proteins identified across bacterial genomes. The outer ring indicates bacterial taxonomic groups, with colors corresponding to the legend. Letters indicate phylogenetic clusters examined in (B) and (C). Protein in clades E and F were non-vault like SPFH family members. The position of *P. aeruginosa* PA14 Band7 is indicated. Scale bar represents 0.25 amino acid substitutions per site. **(B)** Genomic neighborhoods of representative vault-like band7 homologs from the indicated phylogenetic clusters. Genes are colored according to their predicted functions, including transcriptional regulation, RNA ligation, RNA cyclization, and nucleotidyltransferase activity. Putative tRNA-like sequences adjacent to *band7* are highlighted in green, with their predicted amino acid identities indicated when available. Blue shading highlights *band7* and its associated tRNA-like region. Gene organization is shown to scale. **(C)** Predicted oligomeric structures of representative vault-like Band7 proteins from the indicated phylogenetic clusters. An individual Band7 protomer is highlighted in each structure. Structures are colored according to their corresponding taxonomic groups in (A).

## Discussion

Here, we establish that the *P. aeruginosa* RNA repair system mitigates damage by ribotoxins deployed during intermicrobial antagonism. Following intoxication, the RNA cyclase RtcA amplifies signals of RNA damage by producing the chemical signal that activates the transcriptional activator RtcR. RtcR drives expression of the divergently encoded Rtc operon, including genes for: Band7, an uncharacterized protein predicted to assemble into complex homologous to the mammalian vault; RtcB, an RNA ligase; RlaP, a poorly characterized nucleotidyltransferase, and RtcA. Upstream of the Rtc operon, RtcR also activates transcription of bRNA, a noncoding RNA with tRNA-like features. The diversity of Rtc operon constituents across bacteria suggests that these genes have evolved to be coordinately regulated in response to a common hallmark of RNA damage, 2′,3′-cyclic phosphate ends, while individual genes counter distinct forms of RNA damage. Here, we demonstrate roles for Band7, RtcB, and RtcA in countering ribotoxins, however, our panel of toxins did not reveal a protective role for RlaP. Recent work suggested that RlaP counters ribotoxins that cleave RNA and remove one or more nucleotides, preventing repair by RtcB alone (*34*). Testing a wider range of ribotoxins in future studies may capture the specific utility of this gene.

Our findings place Rtc operons within a growing repertoire of genes that protect bacteria against toxins deployed during microbial conflict. These defenses range from very specific, such as acquiring cognate immunity proteins, to very generalized defenses, such as protective barriers that block intoxication (*10*, *35*). Rtc operons are induced by damage inflicted by antibacterial toxins and join a larger family of damage induced responses. A conceptually similar screen for resistance to T6SS toxins in *P. aeruginosa* identified three discrete gene clusters, termed antagonism resistance clusters (Arc1–3), that protect against antibacterial effectors that damage the cell membrane (*36*). Arc1–3 are regulated by the Gac/Rsm danger-sensing system and recently *rtcB* was also shown to be regulated by Gac/Rsm in PAO1 (*37*). This regulatory overlap supports the notion that defense against antibacterial toxins is part of a broad immune response to microbial antagonism and may constitute a new arm of the bacterial immune system. Indeed, the apparent escalation of offensive ribotoxins and the corresponding repertoire of genes within RNA defense systems is reminiscent of the molecular arms race between phages and phage defense systems. This analogy implies that, like phage defense systems, there may be many additional antibacterial toxin defense systems waiting to be discovered.

The most surprising finding of our study is that bacteria encode proteins homologous to the eukaryotic MVP that mitigate RNA damage to allow bacteria to survive treatment with a ribotoxin. Multiple publications have identified MVP homologs (*23*, *38–43*) and two recent studies demonstrated that related Band7 proteins assemble into vault complexes *in vitro* and oligomerize with 70 or more protomers to form particles that rival the human vault in size (*44*, *45*). Although our structural predictions were limited to 12 Band7 protomers, we anticipate PA14 Band7 proteins may also form particles similar in size to human vaults. An outstanding question is, why does ribotoxin defense require a molecular vessel like the vault? We envision at least three potential models. First, bacterial vaults may trap ribotoxins to sequester these dangerous molecules away from cellular tRNAs. Second, the vault may be a cryptic enzyme or may capture another cellular enzyme that requires compartmentalization for specificity or activity. Third, the vault may encapsulate damaged tRNA molecules to facilitate their repair or destruction.

The proposed models for bacterial vaults are not mutually exclusive, however, each final mechanism must have a role for bRNA because this component was necessary for ribotoxin defense. We hypothesize that bRNA serves as the “canary in the coal mine” for toxin presence. By resembling tRNA, the bRNA acts as a decoy substrate for ribotoxins and bRNA cleavage likely activates defense via Band7. However, it cannot be as simple as bRNA holding Band7 in an inactive state until cleavage because expression of Band7 in the absence of bRNA did not result in a constitutive defensive state. Perhaps the 5′ and 3′ extensions that separate bRNA from similar arginyl-tRNAs serve a crucial function for defense activation. Looking past *P. aeruginosa*, the bRNA genes we identify across bacteria are not just similar to arginyl-tRNA but resemble a wide range of other tRNA species. Some even encode multiple tRNA annotations. These observations suggest that each bRNA/Band7 pair may be specific for toxins that cleave different target tRNA. In this way, bRNA must retain the tRNA identity elements recognized by the ribotoxin.

The eukaryotic vault complex has mystified researchers for over 40 years (*46*). While this nanoparticle has been implicated in diverse cellular processes and disease states, a molecular explanation for how the vault contributes to cellular homeostasis has yet to emerge. A unique challenge of understanding the vault has been the absence of these complexes in conventional eukaryotic model organisms such as *Saccharomyces cerevisiae*, *Caenorhabditis elegans*, and *Drosophila melanogaster* (*47*). Our findings establish that bacteria could be the missing model organism for these observations. Accordingly, we hypothesize that the human vault complex may play a crucial part in repairing or responding to RNA damage.

## Acknowledgements

The authors would like to thank the CU Boulder Department of Biochemistry Shared Instruments Pool core facility (RRID:SCR_018986), Anette Erbse, and its staff; Karolin Luger and her lab for sharing lab space, resources and equipment; Michael Nosalla for helpful discussions; John Whitney for strains and plasmids; and members of the Whiteley laboratory, particularly Kishen Patel, for their advice, support, and helpful discussions.

## Authors Contributions

Conceptualization: NPB and ATW

Methodology: NPB and ATW

Investigation: NPB, LCH, LKR

Visualization: NPB and ATW

Funding acquisition: ATW

Supervision: ATW

Writing - original draft: NPB and ATW

Writing - review & editing: NPB, TAN, and ATW

## Data, code, and material availability

All data are available in the manuscript or supplementary materials. No new code was generated for this study. Strains and plasmids used in this study are available following publication upon request and/or are distributed by Addgene (addgene.org).

## List of Supplementary Materials

Tables S1 to S2

## Materials and Methods

### Bacterial strains and bacteriophages

Bacteria used in this study are listed in Supplementary Table 1. All strains were cultured in LB medium (1% tryptone, 0.5% yeast extract, and 0.5% NaCl) at 37 °C with shaking at 220 rpm in 1-3 mL of medium in 14 mL culture tubes, unless otherwise indicated. Where applicable, cultures were supplemented with gentamicin (15 µg/mL for *E. coli* and 30 µg/mL for *P. aeruginosa*) or carbenicillin (100 µg/mL). Bacterial strains were frozen for storage in LB supplemented with 10% glycerol at -70 °C*. E .coli* Omnipir was used for the construction and propagation of all plasmids*. E. coli* SM10 was used for mating the allelic exchange vector pEXG2 into all *P. aeruginosa* strains, and *E. coli* Rosetta2 was used for all heterologous recombinant protein expression. *E. coli* MG1655 and *P. aeruginosa* PA14 were used to collect all experimental data. When growing strains that required induction, 250 µM or 500 µM β-D-1-thiogalactopyranoside (IPTG) were used to induce, as appropriate.

All bacteriophages used in this study are listed in Supplementary Table 1.

### Cloning, plasmid construction and mutant strain generation

Plasmids used in this study are listed in Supplementary Table 3. Genes of interest were amplified from bacterial genomic DNA using Q5 Hot Start High Fidelity Master Mix (NEB M0494L), flanked by 18 base pairs of homology to vector backbone. Amplified genes were ligated into restriction-digested, linearized vector backbones using Gibson assembly(*48*). Gibson reactions were then transformed into bacteria by electroporation and plated on the appropriate antibiotic selection. All enzymes used for cloning were purchased from New England Biolabs.

Mutants of *P. aeruginosa* were generated through allelic exchange as previously described (*49*). Mutants of *E. coli* were generated through Lambda red recombineering as described previously (*50*).

### Recombinant protein expression and purification

For protein expression and purification of pyocin S4 and the S4 chimeras, each toxin was co-expressed with its cognate immunity protein from a single pETDuet vector to prevent cellular intoxication and to increase yield of recombinant protein. 2 L of LB broth supplemented with carbenicillin was inoculated 1:50 with an overnight culture of *E. coli* Rosetta2 harboring the toxin-immunity construct. Cultures were grown at 37°C with shaking at 220rpm until they reached an OD_600_ of 0.4, and IPTG was added at a final concentration of 500 µM IPTG prior to growth for an additional 3 h. Cells were harvested by centrifugation at 4,660 x *g* and 4°C, pellets were then resuspended in lysis buffer (100 mM Tris-HCl, pH 8.0, 300 mM NaCl, 10 mM imidazole), and sonicated with a Sonicator 4000 (Misonix) on ice at an amplitude of 75% for six 30 second on / off pulses. Lysates were cleared of insoluble debris by centrifugation at 39,000 x *g* for 60 min. Soluble fractions were then loaded onto 1.5-mL gravity-flow columns packed with 1 mL of Ni-NTA agarose, pre-equilibrated with lysis buffer. The column-bound toxin-immunity complexes were washed three times with 20 mL of lysis buffer and subsequently eluted using 5 mL of lysis buffer supplemented with 400 mM imidazole. Proteins in each Ni-NTA elution were then further purified by fast protein liquid chromatography using gel filtration on a HiLoad 16/600 Superdex 200 column equilibrated in 20 mM Tris-HCl, pH 8.0, and 150 mM NaCl. The purity of each protein was determined by SDS-PAGE followed by gel staining with Coomassie Brilliant Blue.

### SDS-polyacrylamide gel electrophoresis

Purified pyocin S2, S4 and the S4 chimeric toxins were diluted 1:2 in 4X Laemmli buffer supplemented with 1.5% β-mercaptoethanol. Samples were heated to 95 °C for 10 minutes and then loaded on a SurePage^TM^, Bis-Tris 4-20% gradient polyacrylamide gel (Genscript). Protein samples were resolved for 50 minutes at 155V before being stained with Coomassie brilliant blue and visualized using a Azure 200 Gel Imaging Workstation (Azure Biosystems).

### Bacterial intoxication spot assays

Spot assays were used to measure the sensitivity of *P. aeruginosa* strains to pyocin S2, S4 and the S4-chimera ribotoxins. Overnight cultures of *P. aeruginosa* were back diluted 1:50 in LB and cultivated at 37 °C with shaking at 220 rpm to a final OD_600_ of 0.4. Cultures were serially diluted 10-fold into fresh LB. From each 10-fold dilution (ranging from 10^-1^ to 10^-8^), 10 µL of culture was added to the well of a microtiter plate containing 87 µL of LB and 3 µL of toxin, and the resulting mix spotted onto LB agar plates. Plates were incubated overnight at 37 °C prior to imaging.

The amount of toxin needed for measurable intoxication was empirically determined for each toxin. Final concentrations of toxin used in Fig. 2D were 80 nM for S4-EndoU^tox^, 50 nM for S4-S4^tox^, 69 nM for S4-D1^tox^, 900 nM for S4-D2^tox^, 690 nM for S4-S6^tox^, and 89 nM for S2. For all other spot assay intoxications, these values were increased approximately 10-fold to better measure the relative growth inhibition between mutants and wildtype *P. aeruginosa.*.

### Temperature Stress Assay

Overnight cultures were diluted 1:50 into fresh LB medium and cultivated at 37 °C with shaking at 220 rpm to an OD600 of 0.5. Cultures were serially diluted and spotted onto LB agar plates. Plates were incubated at 10 °C, 22 °C (room temperature), or 37 °C. Plates incubated at 22 °C and 37 °C were imaged following overnight growth, whereas plates incubated at 10 °C were imaged after 3 days, when visible colonies had developed.

### UV exposure assays

Overnight cultures were back diluted 1:50 in LB and cultivated to an OD_600_ of 0.4. 100 µL of cells were pelleted by centrifugation at 8,000 x *g* and resuspended in 100 µL of phosphate buffered saline (PBS,137 mM NaCl, 2.7 mM KCl, 10 mM Na_2_HPO_4_, and 1.8 mM KH_2_PO_4_, pH 7.4). 50 µL of the resuspended pellet was then spotted onto the lid of a petri dish and placed drop side up into a Stratalinker UV crosslinker (Stratagene) operating at 254 nm. Samples were exposed to UV irradiation (80-120 µJ/cm^2^), serially diluted 10-fold in PBS, then spotted on 1.6 % LB agar plates and grown overnight at 37 °C for quantification of cell viability.

### Disk diffusion assays

Overnight cultures were diluted 1:50 into fresh LB and cultivated to an OD_600_ of 0.4. 300 µL of culture was then mixed with 3.5 mL of 0.35% LB agarose warmed to 40 °C. This mixture was then poured onto a 1.6 % agar LB plate and cooled for 10 minutes to solidify. One cm diameter disks of filter paper were soaked with 50 µL of streptomycin (100 µg/mL), kanamycin (50 µg/mL), gentamicin (30 µg/mL), spectinomycin (100 µg/mL), apramycin (15 µg/mL), chloramphenicol (20 µg/mL), tetracycline (15 µg/mL), carbenicillin (100 µg/mL), polymyxin b (30 µg/mL), mitomycin (5 µg/mL), zeocin (50 µg/mL), ciprofloxacin (5 µg/mL), rifampin (5 µg/mL), irgasan (24 µg/mL), or H_2_O_2_ (3% solution). Impregnated disks were then placed onto the cooled agar overlay and plates were incubated overnight at 37 °C.

### Efficiency of plating/phage replication analysis

A modified double agar overlay assay was used to measure phage replication. Briefly, overnight cultures of the indicated strains were diluted 1:100 and cultivated for an additional two hours to reach logarithmic growth phase (OD_600_ 0.1-0.8). 400 µL of the culture was then mixed with 3.5 mL 0.35% LB agar, plus an additional 10 mM MgCl_2_,10 mM CaCl_2_, and 0.1 mM MnCl_2_. The mixture was then poured onto a 1.6% agar LB plate and cooled for 15 minutes. 2 µL of a phage dilution series in SM buffer (100 mM NaCl, 8 mM MgSO_4_, 50 mM Tris-HCl pH 7.5, 0.01% gelatin) was spotted onto the overlay and allowed to dry under a flame before incubation overnight at 37 °C. Plaque formation was analyzed the following day. When no plaques were observable or a hazy zone of clearance occurred, 0.9 plaques at the least dilute spot was used as the limit of detection.

### Intoxication of bacterial cells for RNA extraction

For isolation of RNA from intoxicated cells for RNA sequencing or Urea-PAGE, overnight cultures of *P. aeruginosa* PA14 were back diluted 1:50 and cultivated to an OD_600_ 0.4. 500 µL of culture was then mixed with 20 µL of purified S4-S4EndoU^tox^ for a final concentration of 1.6 µM, and incubated at room temperature for 5, 15, 30, 45, or 60 minutes. Bacterial cells were then pelleted by centrifugation at 8,000 x *g* and frozen at -70 prior to RNA extraction and purification.

### Extraction, purification and sequencing of RNA

Cell pellets were resuspended in 1 mL of TRIzol reagent (ThermoFisher), vortexed for 10 seconds, and incubated at room temperature for 5 minutes. Chloroform (200 µL) was added, and samples were vortexed for 30 seconds before incubation on ice for 2 minutes. Samples were briefly mixed and centrifuged at 14,000 x g for 15 minutes at 4 °C to separate the aqueous and organic phases. Approximately 500 µL of the upper aqueous phase was transferred to an equal volume of ice-cold isopropanol, vortexed for 15 seconds, and incubated on ice for 20 minutes. RNA was pelleted by centrifugation at 14,000 x g for 15 minutes at 4 °C, washed three times with 1 mL of 75% ethanol, and air-dried. Pellets were resuspended in 100 µL of DNase I reaction buffer (NEB; 10 mM Tris-HCl pH 7.6, 2.5 mM MgCl2, 0.5 mM CaCl2) and treated with 4 units of DNase I (NEB) for 30 minutes at 37 °C. RNA was further purified using the Monarch RNA Cleanup Kit (NEB) according to the manufacturer’s instructions and eluted in DEPC-treated water. RNA concentration and purity were assessed via absorbance at 254 nm using a NanoDrop One^C^. Sample concentrations were adjusted to a final concentration of 100 ng/µL and stored at -80 °C. RNA was sequenced using Illumina with rRNA depletion and number of reads (SeqCenter).

### RNA-seq read processing and alignment

Paired-end sequencing reads were quality-filtered and trimmed using fastp before alignment to the *P. aeruginosa* PA14 reference genome (NC_008463) using Bowtie2 with the --very-sensitive and --end-to-end settings (*51*, *52*). Resulting alignments were sorted and indexed using SAMtools (*53*). Gene-level read counts were generated using featureCounts with reverse-stranded, paired-end fragment counting (-s 2 -p --countReadPairs), using the NCBI PA14 genome annotation and grouping features by locus tag (*54*).

### Transcript abundance and expression analysis

Transcript abundance was calculated as transcripts per million (TPM) by normalizing gene-level fragment counts to annotated gene lengths and subsequently scaling to one million transcripts per sample. Relative expression was determined by comparing TPM values between all experimental conditions (EndoU^tox^ exposure for 5 and 15 minutes, and heterologous expression of RtcR^ΔCARF^) and the empty vector control (no toxin, no induction). Genes exhibiting at least a 2.5-fold change in either direction, with a minimum abundance of 10 TPM in at least one condition, were classified as differentially expression. A pseudocount of 1 TPM was applied when calculating fold changes. Coverage tracks were generated using deepTools bamCoverage with counts-per-million (CPM) normalization and a bin size of 1 nucleotide (*55*). Coverage tracks were visualized using Integrative Genomics Viewer (IGV) (*56*).

### Urea-polyacrylamide gel electrophoresis

200 ng of purified RNA in water was diluted 1:2 with 2x RNA loading dye (NEB). Sample were heated to 80 °C for 2 minutes to denature RNA and then placed on ice for 5 minutes. Samples were then separated using 10 cm TBE-denaturing acrylamide gels (8.25% v/v acrylamide, 8M urea,1X TBE) at 200V for 50 minutes in 1X TBE buffer. Gels were stained for 10 minutes with SYRB gold (ThermoFisher) and visualized using an Azure 200 Gel Imaging Workstation (Azure Biosystems).

### Northern Blotting

RNA resolved by urea-PAGE was transferred to a positively charged quaternary amine nylon membrane (Roche) via wet transfer at 0.4 A for 45 minutes using a BioRad Mini Trans-Blot electrophoretic transfer system. RNA was crosslinked to the membrane using a Stratalinker UV crosslinker with the “Auto Crosslink” setting, and the membrane was subsequently transferred directly into 10 mL of ULTRAhyb-Oligo hybridization buffer (Thermo Fisher Scientific), preheated to 42 °C in a shaking incubator rotating at 70 rpm. After prehybridization for 10 minutes, biotinylated DNA probe was added to the buffer at a final concentration of 3 nM. Hybridization was performed overnight, and membranes were washed twice for 10 minutes each with 10 mL of stringency wash buffer (2x SSC, 5% SDS, w/v) at 42 °C with gentle agitation. Blots were then moved to room temperature and washed twice for 10 minutes with 10 mL of wash buffer (PBS, pH 7.4, 0.5% SDS, w/v), followed by three washes with 10 mL of blocking buffer (PBS, pH 7.4, 0.5% SDS, w/v, 0.1% Thermo Fisher I-Block reagent). Blots were left in blocking buffer for 1 hour at room temperature. Licor IRDye 800CW was diluted 1:5,000 into 12 mL of fresh blocking buffer and added to the blot. After 1 hour of incubation with gentle agitation, the blot was washed twice with 10 mL of blocking buffer and then twice with 10 mL of wash buffer. Blots were subsequently imaged using a Licor Odyssey CLx imaging system. Following imaging, RNA ladder positions on Northern blots were determined by overlaying an image of the acrylamide gel containing separated RNA stained with SYBR Gold prior to transfer and marking the corresponding positions of the RNA standards on the blot.

### Sequence database and profile HMM search

The Band7 protein from *P. aeruginosa* PA14 was used as a query in iterative PSI-BLAST searches (2 rounds) to identify 19 candidate vault-like Band7 proteins from diverse bacterial species. Candidate proteins were manually inspected using AlphaFold structural models available through InterPro to confirm the presence of N-terminal globular repeat domains characteristic of vault proteins. The 19 sequences were aligned using MAFFT, and the resulting alignment was used to construct a profile Hidden Markov model (HMM) using HMMER hmmbuild (*57*, *58*). The HMM was searched against a local database of approximately 88.8 million nonredundant protein sequences derived from bacterial and archaeal NCBI RefSeq assemblies using hmmsearch. Hits were retained using a full-sequence E-value threshold of ≤1 x 10^-30^, a minimum protein length of 150 amino acids, and exclusion of proteins with NP_ or YP_ accession prefixes, yielding 2,026 candidate vault-like Band7 proteins.

### Phylogenetic and gene neighborhood analyses

Candidate proteins were clustered using MMseqs2 at 90% sequence identity and 80% alignment coverage, yielding 1,312 representative proteins (*58*). Taxonomic assignments were obtained using NCBI taxonomy, and archaeal sequences and proteins with unresolved bacterial taxonomic assignments were excluded. The remaining representative proteins and 19 initial query sequences were aligned using MAFFT (--auto) (*58*). An approximately maximum-likelihood phylogeny was inferred using FastTree with default parameters and visualized using Interactive Tree of Life (iTOL) (*59*, *60*).

### Genomic neighborhood analysis of vault-like Band7 proteins

The genomic neighborhoods of Band7 homologs were examined using FlaGs2 (Flanking Genes) (*61*). RefSeq protein accession numbers were used to retrieve genomic regions encompassing five protein-coding genes upstream and downstream of each Band7 homolog (-g 5). Gene neighborhoods were compared across bacterial taxonomic groups to identify conserved gene associations. Genomic regions were retrieved from NCBI as annotated GenBank files and visualized using Geneious Prime version X.

### Structural modeling of bacterial vault-like Band7 proteins

Protein structures of bacterial vault-like Band7 homologs were predicted using the AlphaFold online server (*62*). Predicted structures were visually inspected to assess conservation of the vault-like architecture, including variation in the number of N-terminal repeat domains and the length of the C-terminal cap helix. Structural models were visualized and rendered using UCSF ChimeraX(*63*).

### Multiple sequence alignment of P. aeruginosa arginyl-tRNA sand bRNA

Annotated *P. aeruginosa* PA14 arginine tRNA sequences and the tRNA-like region of bRNA (PA14_60630) were extracted and aligned using the Geneious multiple sequence alignment tool with default settings.

### Secondary structure prediction of arginyl-tRNA and bRNA

The mimimum free energy secondary structures of *P. aeruginosa* arginyl-tRNA^ACG^ and tRNA-like region of bRNA was modelled using the Vienna RNAfold webserver under default settings (*64*).

## Supplementary Figures

**Figure S1.**
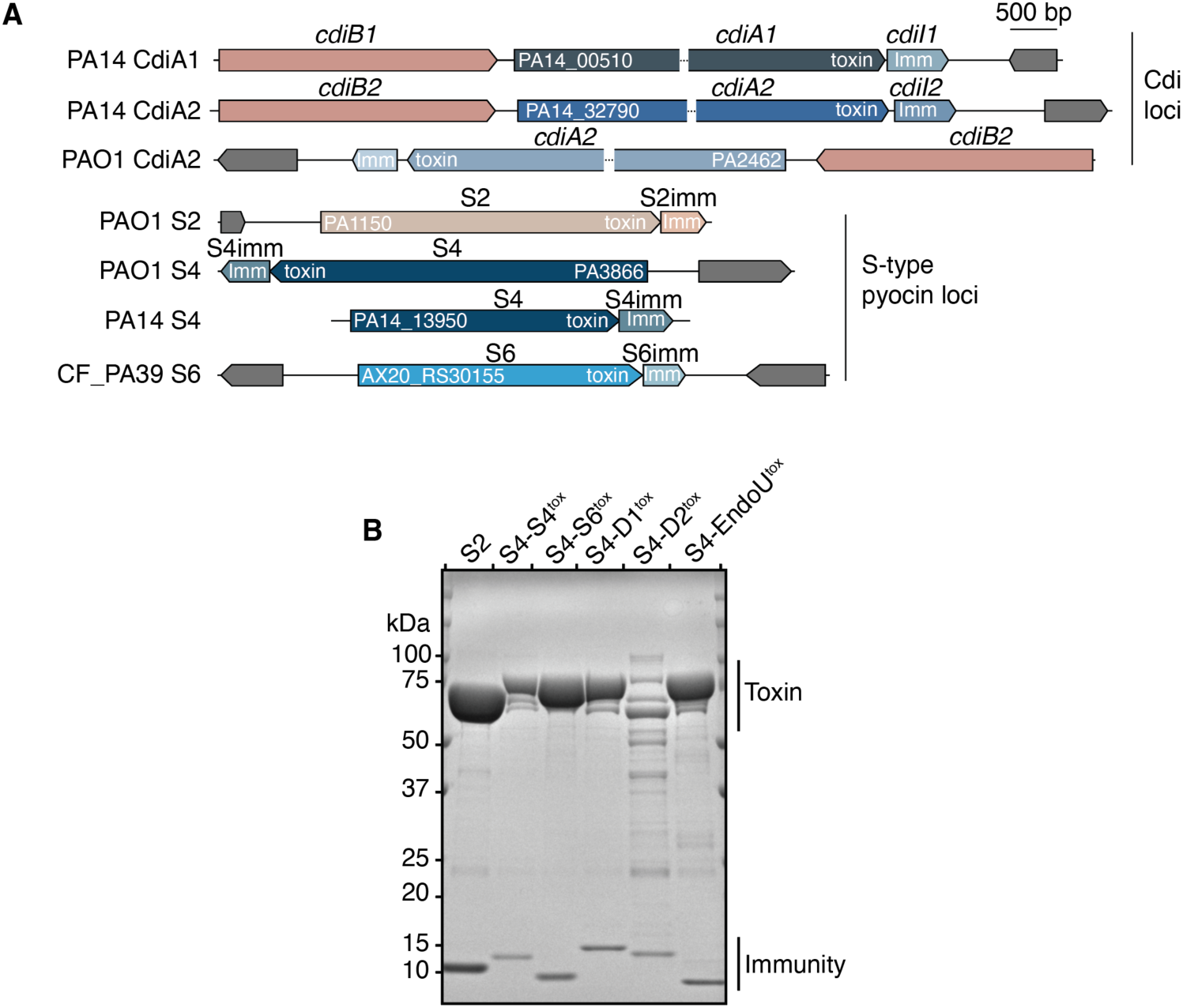
Generation of chimeric ribotoxins. **(A)** Genomic organization of the Cdi and S-type pyocin loci encoding toxins used to generate the engineered ribotoxin panel, together with the DNA-targeting pyocin S2 used as a control. Toxin domains and cognate immunity genes labeled “toxin” and “Imm,” respectively. The bacterial strains from which each locus originates are indicated. Breaks in *cdiA* genes indicate regions of DNA omitted for clarity. **(B)** Polyacrylamide gel electrophoresis of purified pyocins S2, S4, and chimeric S4 toxins co-expressed with their respective immunity proteins. The predicted molecular weights of the toxin and immunity proteins, respectively, are as follows: S2 (74 and 10 kDa), S4-S4^tox^ (81 and 13 kDa), S4-S6^tox^ (81 and 9 kDa), S4-D1^tox^ (83 and 18 kDa), S4-D2^tox^ (84 and 18 kDa), and S4-EndoU^tox^ (85 and 10 kDa).

**Figure S2.**
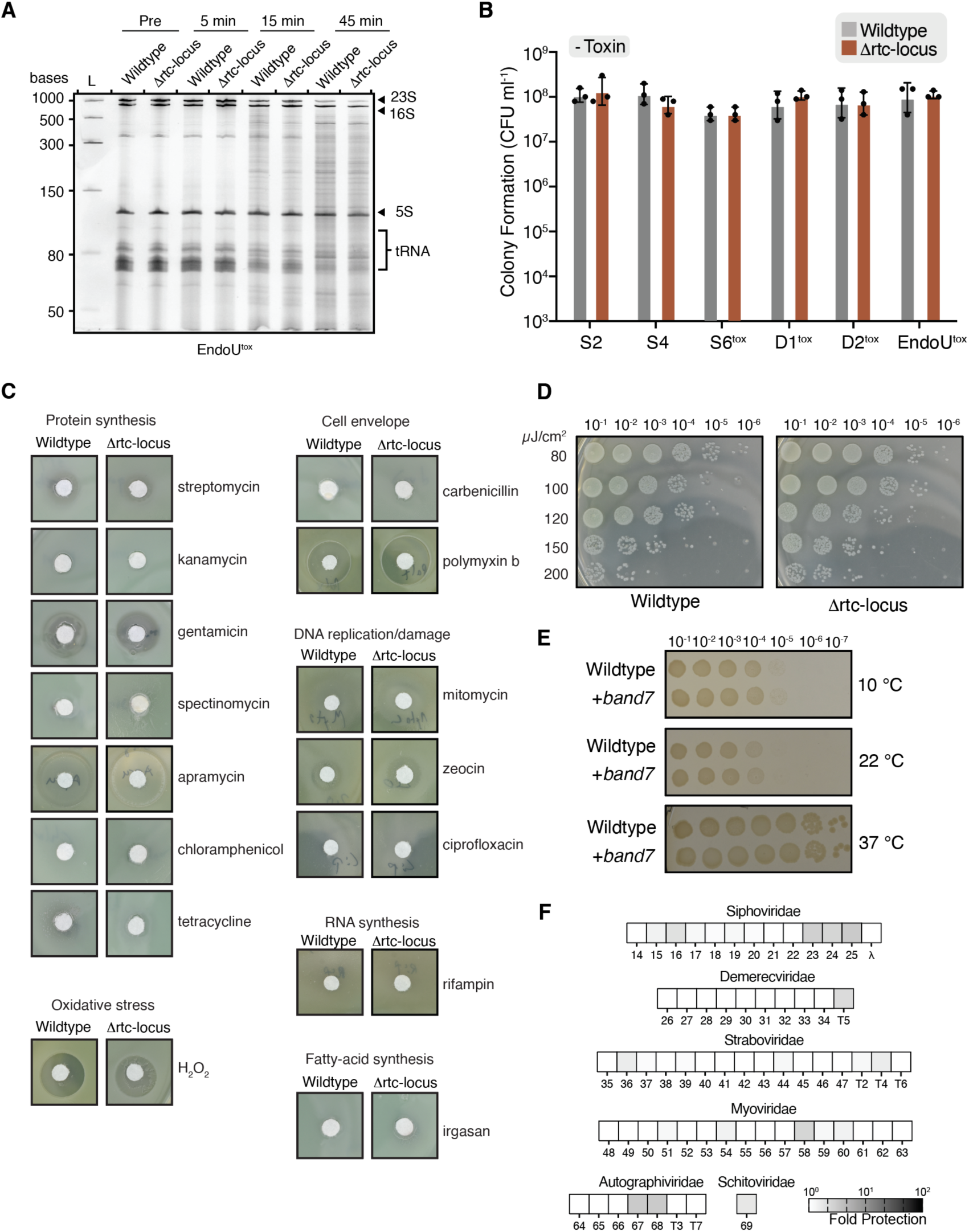
Challenge of the Δrtc-locus mutant with diverse stressors. **(A)** Urea-PAGE analysis of total cellular RNA extracted from *P. aeruginosa* exposed to S4-EndoU^tox^ for the indicated time points. Major RNA species and cleavage products are indicated. **(B)** Quantification of colony-forming units (CFU) of wildtype and rtc-locus *P. aeruginosa* in the absence of toxin. Data are the mean ± SEM of three biological replicates (n = 3), with individual replicates shown as points. **(C)** Disk diffusion assays comparing sensitivity of wildtype and Δrtc-locus *P. aeruginosa* to the indicated antibiotics and other stressors. Compounds are grouped according to the cellular targets or processes they inhibit. **(D)** UV sensitivity of wildtype *P. aeruginosa* and the Δrtc-locus mutant. Cultures were serially diluted tenfold and spotted onto LB agar before exposure to increasing doses of UV irradiation (80-200 μJ/cm^2^). **(E)** Temperature sensitivity of wildtype *E. coli* MG1655 and *E. coli* MG1655 carrying the Band7-containing locus from *E. coli* ECOR67. Cultures were serially diluted tenfold, spotted onto LB agar, and incubated at 10 °C, 22°C, or 37 °C. The Ecor67 Band7 protein is a homolog to PA14 Band7. **(F)** Plaque formation (PFU mL^-1^) was enumerated for each phage infecting wildtype *E. coli* MG1655 or MG1655 carrying the chromosomally integrated Band7-containing locus from *E. coli* ECOR67. The Ecor67 Bandy protein is a homolog to PA14 Band7. Fold protection was calculated by dividing the phage titer of wildtype MG1655 by that on MG1655 carrying the ECOR67 locus and depicted as a heatmap. Phages are grouped by family, indicated above each group, with numbers corresponding to their identifiers in the BASEL phage collection(*65*).

**Figure S3.**
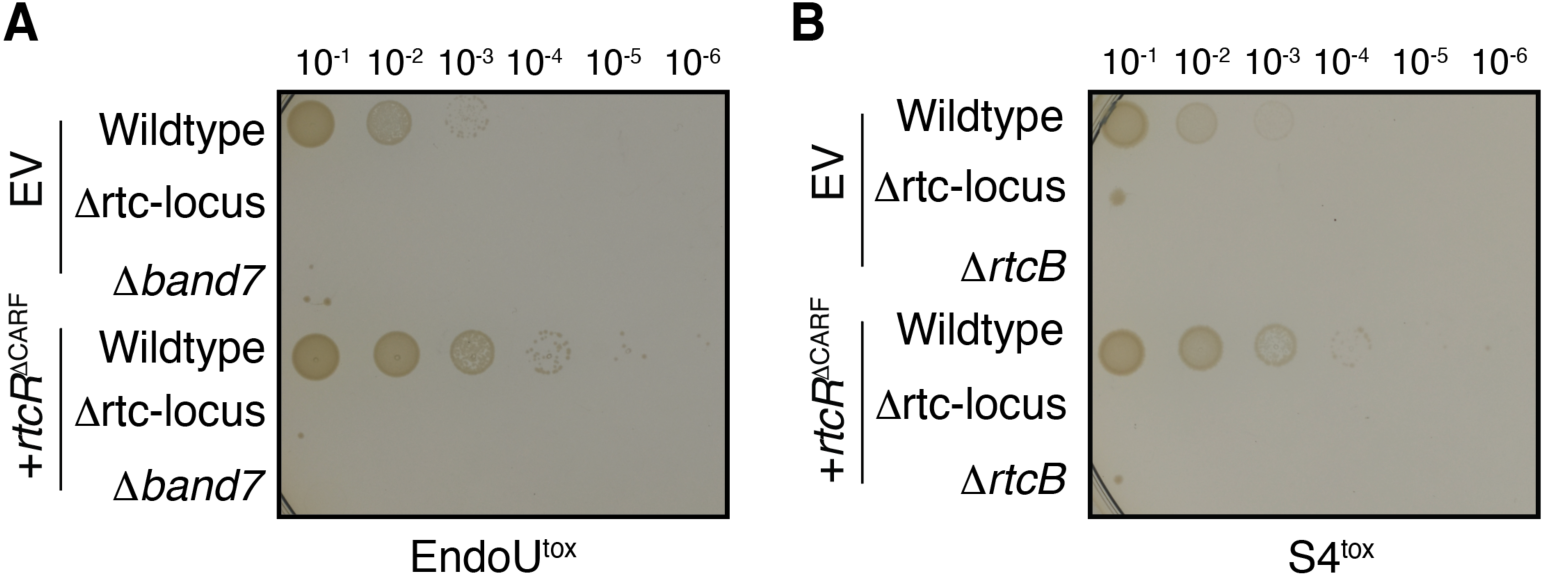
Constitutively active RtcR^ΔCARF^ does not complement the Δ*band7* or Δ*rtcB* mutants. **(A and B)** Toxicity assays showing tenfold serial dilutions of the indicated *P. aeruginosa* strains expressing *rtcR*^ΔCARF^ or carrying an empty vector control (EV), following exposure to S4-EndoU^tox^ (A) or S4-S4^tox^ (B). Cells were spotted onto LB agar supplemented with 250 μM IPTG to induce protein expression.

**Figure S4.**
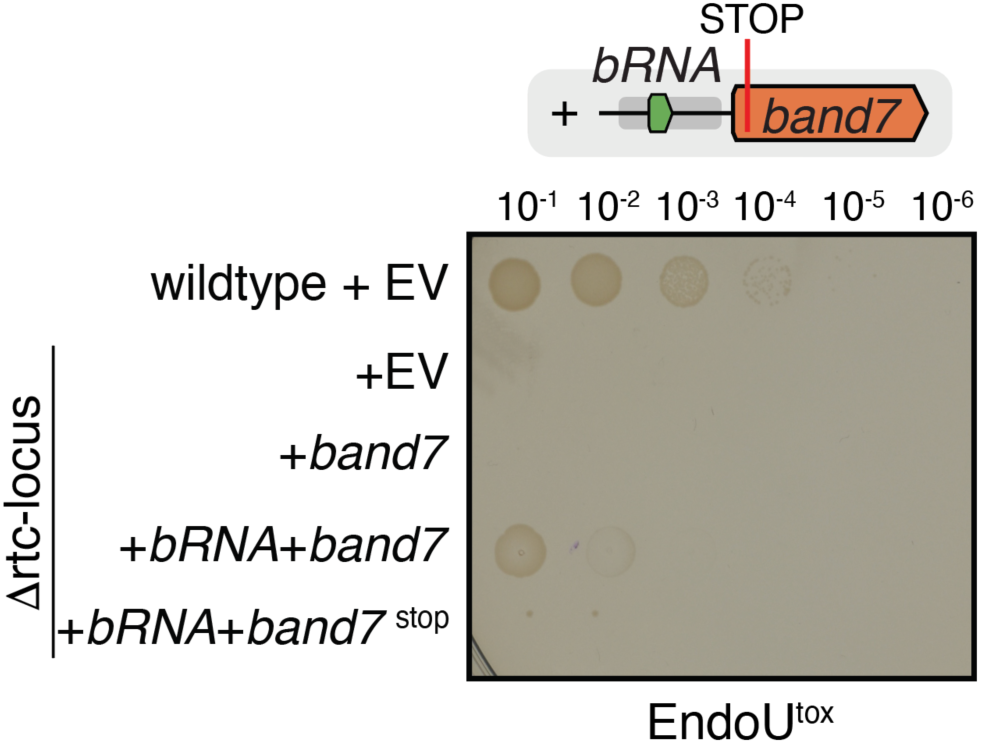
Requirement of Band7 and bRNA for protection against ribotoxic stress. **(A)** Tenfold serial dilutions of wildtype *P. aeruginosa* or the Δrtc-locus strain carrying either an empty vector (EV) or the indicated construct, following intoxication with S4-EndoU^tox^. Cells were spotted onto LB agar supplemented with 250 μM IPTG. Introduction of a premature stop codon in the *band7* reading frame abolished protection against S4-EndoU^tox^.

**Figure S5.**
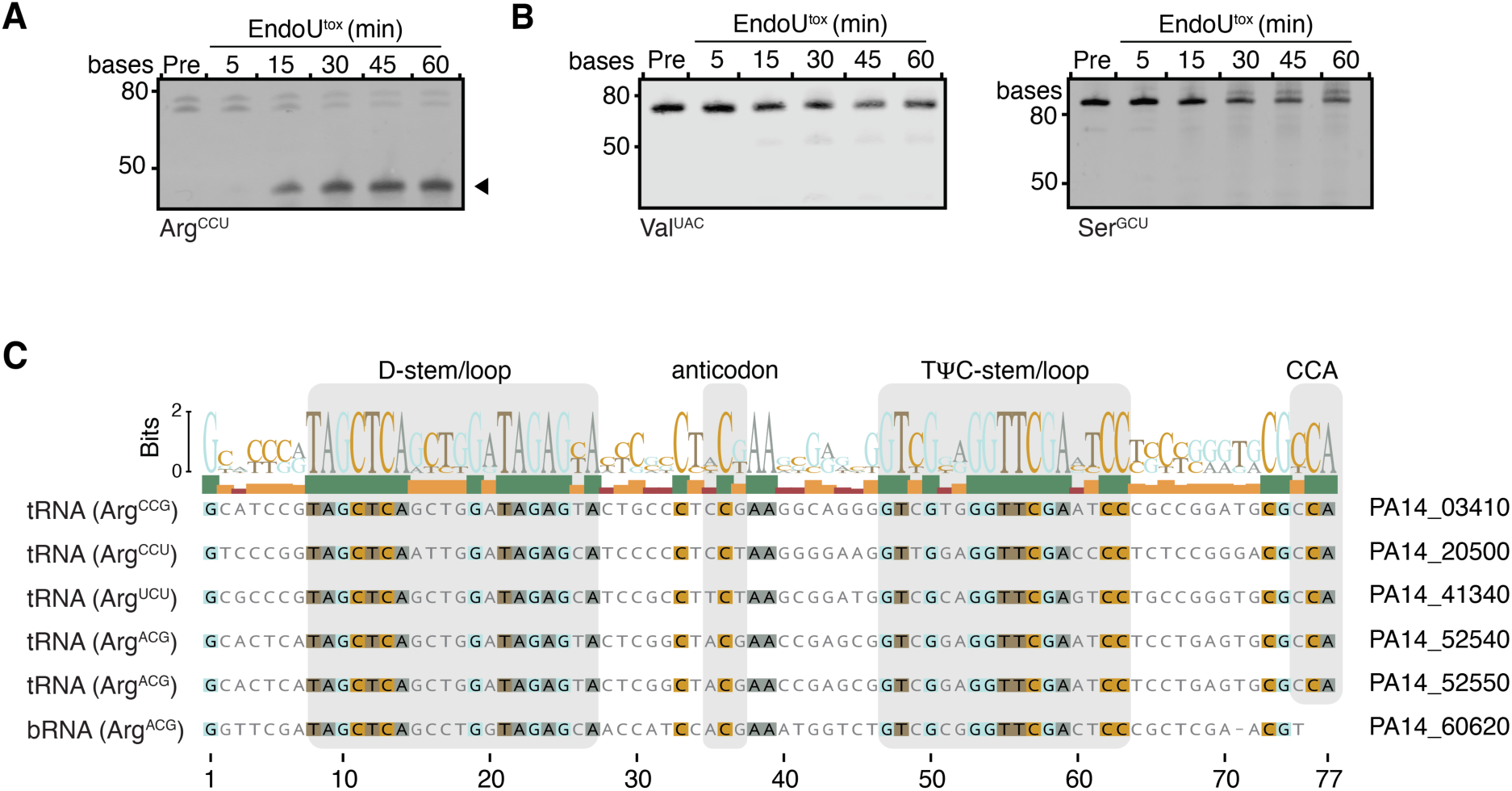
bRNA retains conserved features of tRNAs. **(A)** Northern blot analysis of tRNA Arg^CCU^ cleavage following intoxication with EndoU^tox^. RNA was isolated from *P. aeruginosa* before intoxication (Pre) or at the indicated times post-intoxication. Arrowheads indicate major cleavage products. **(B)** Northern blot analysis of tRNAs Arg^CCU^ and Ser^GCU^ following intoxication with EndoU^tox^. RNA was isolated from *P. aeruginosa* before intoxication (Pre) or at the indicated times post-intoxication. **(C)** Multiple sequence alignment of five annotated arginine tRNAs in *P. aeruginosa* and the 74-nucleotide tRNA-like region of bRNA (PA14_60620). Conserved sequence motifs corresponding to the D-stem/loop, anticodon region, and the T-stem/loop are highlighted. The sequence logo above the alignment indicates nucleotide conservation at each position.

## Supplementary Tables

**Table S1. Toxins used by *P. aeruginosa* during intermicrobial warfare**

A list of antibacterial toxins produced by *P. aeruginosa* with characterized bacterial killing activity. A non-exhaustive literature search identified 52 antibacterial toxins deployed through type six secretion, contact-dependent inhibition, or pyocin release. Note that that antibacterial activity has not been directly demonstrated for the CdiA1 and CdiA2 loci in *P. aeruginosa* PA14. However, as the equivalent loci in strain PAO1 have been characterized as having antibacterial activity, we included them in this list. We demonstrate their antibacterial properties in this study. References to studies showing antibacterial activity are included as Pubmed IDs under the row titled “Reference (Pubmed ID)”.

**Table S2. Raw read counts and transcripts per million (TPM) values from RNA sequencing.**

